# Lipid Anchor Chemistry and Spacer Architecture Influence DNA Availability and Light-Directed Cell Printing

**DOI:** 10.64898/2026.11.10.765690

**Authors:** Danielle Herubin, Katelyn Mathis, Lamees Abu-Suleiman, Jenna Li, Brian Meckes

**Affiliations:** Department of Biomedical Engineering, University of North Texas, 3940 N Elm St., Denton, TX 76207, USA; BioDiscovery Institute, University of North Texas, 1155 Union Circle Denton, Texas 76203; Texas Academy of Math and Science, University of North Texas, 1155 Union Circle Denton, Texas 76203

**Keywords:** Cell printing, DNA assembly of cells, light-directed bioprinting

## Abstract

Precise control over cellular interactions is critical to understanding the complex underpinnings of disease. DNA-printing of cells (DPC), a technology that uses complementary DNA to program cellular interactions, has the potential to enable precision cell printing. However, the simplicity of printing is undermined by the stability of anchoring moieties. This manuscript systematically investigates how commercially available hydrophobic DNA anchor and co-anchor chemistries influence DNA availability and light-directed DPC (LD-DPC) printability. Using cholesterol, tocopherol, and cholesterol-tocopherol modifications with defined spacer architectures, we evaluated how anchor chemistry and molecular architecture influence functional DNA presentation at the cell surface. Notably, incorporating additional polyethylene glycol spacers between lipid anchors and DNA enhances functional DNA availability and patterning performance. While cholesterol-based anchors provide the highest initial surface loading, tocopherol-containing anchors promote greater spatial dispersion and cell-capture efficiency during printing. Notably, multiple anchor/co-anchor combinations maintain functional DNA presentation at room temperature and in serum-containing media. These findings demonstrate that lipid anchor identity and spacer architecture strongly influence the availability of membrane-associated DNA and the performance of light-directed DNA printing of cells. Collectively, these results provide practical guidance for designing stable nucleic-acid-modified cell interfaces using commercially available lipid-DNA chemistries.

## Introduction

Increasingly, biomimetic models that mimic tissues play an important role in modeling disease and development for a myriad of applications including drug development, tissue engineering, and personalized medicine.^1–3^ While many 3D printing technologies are rapidly being developed to meet these needs, DNA printing of cells (DPC) has emerged as a technology that enables systematic layer-by-layer cell placement. DPC leverages the precision of DNA bonding to program cell interactions.^4–9^ Specifically, cells are coated with a single strand of DNA and then are incubated with cells coated with the complementary strand.^4, 6, 10^ These complementary strands then hybridize to force cells to interact. As an extension of this technique, Mathis et al. utilized oligonucleotides with photocages;^11^ the photocages on the bases prevent hybridization until exposed to light.^12–14^ By directing light to specific locations, one can realize light-directed DPC (LD-DPC).^11, 15–17^ Despite the potential power of DPC technologies, stable and affordable cell attachment is critical to their adoption.^4^

Numerous methods for attaching DNA to cell surfaces have been described. As an initial strategy, NHS-ester-terminated DNAs were created to directly attach to amines on the surface of cells.^18, 19^ However, competing hydrolysis reactions during bioconjugation can limit yield, resulting in significant DNA waste.^20^ As a workaround, groups have used two-step amine modification. This strategy involves converting cell-surface amines into clickable tetrazines.^11, 15^ However, this will result in poorly controlled protein modifications; these modifications can persist due to slow protein turnover, which can interfere with protein activity.^21^ A related technique involves the metabolic addition of azide sugars to the glycocalyx or the biotinylation of glycans for bioorthogonal DNA attachment.^22–24^ Again, this requires specialized treatment of cells and extended wait times.^13^

As an alternative approach, lipidated DNAs, i.e., DNAs with hydrophobic terminal modifications, can be incubated with cells and spontaneously inserted into a lipid bilayer. Selden et al. showed that incubation of DNAs terminated in 16 carbon chains allows for semi-stable initial insertion.^10^ To improve stability, two-step processes are required, as it is essential that the DNA does not strongly assemble into micelles while remaining sufficiently strong to remain in the cell membrane.^25–27^ In this process, a longer anchor strand with a locking region and a recognition handle terminated with a hydrophobic anchor is first added to the cell. It is then followed by a shorter co-anchor strand that is complementary to the lock region, terminated with a weaker hydrophobic anchor.^26, 28, 29^ In this fashion, lignoceric acid anchors (24 carbons) with palmitic acid (16 carbons) co-anchors showed greater incorporation efficiency. However, the 24-carbon modifications are not readily available for DNA synthesizers and are costly when performed by vendors. Other groups have pursued cholesterol on both the anchor and co-anchor strands as a more cost-efficient approach.^4^ However, cholesterol-modified oligos are less stable than their lipid chains, losing nearly 25% of their potency^30^ and undergoing endocytosis when incubated in serum.^31^ Printing with cholesterol anchors/co-anchors is performed on ice to limit loss/endocytosis. Facile methods that enable more stable interactions with cell surfaces and greater availability would facilitate cell modification for DPC printing.

Here, we systematically examined how commercially available lipid-DNA anchor and co-anchor architectures influence functional DNA presentation at the cell surface and the resulting performance of light-directed DNA printing of cells. We examined the stability and printability of DNA with cholesterol and tocopherol as anchor/co-anchor strands. Our results demonstrate that adding a polyethylene glycol spacer between the DNA and the hydrophobic anchor improves anchoring stability. When exploring DNA combinations, we found that cholesterol and tocopherol anchor/co-anchor pairs improved printing density. Collectively, this work shows that small modifications to DNA design can enhance LD-DPC printing capabilities by improving retention of accessible DNA at the cell surface. Importantly, this study is motivated by the broader need to understand how lipid identity and molecular flexibility govern the persistence and lateral availability of functional molecules at living cell–material interfaces, rather than by optimization of a single printing workflow.

## Results and Discussion

### Evaluating the Design of DNA

As our first objective, we sought to determine how lipid-anchor chemistry and molecular architecture influence the availability of membrane-anchored DNA at the cell-material interface. We hypothesized that DNA could become unavailable from the cell surface through three mechanisms: 1) dissociation of lipidated DNA from the cell surface membrane, 2) endocytosis of lipidated DNA into the cells, or 3) degradation of DNA on the cell surface. To limit degradation, we incorporated two phosphorothioate linkages at both ends of the DNA, which are known to significantly reduce susceptibility to nucleases. Thus, we focused on changes in surface-associated DNA that could arise from these mechanisms.

We first sought to evaluate how to reduce effects that promote dissociation from the surface. Recent research has shown that the close proximity of double cholesterol anchors on a single DNA promotes dissociation from the surface. Keeping cholesterol far apart on the cell surface improves the stability of lipid membranes.^32^ Current anchor/co-anchor strand designs consist of an anchor srtand terminated with a cholesterol, with one region near the cholesterol complementary to the co-anchor and a secondary capture region for connecting cells **(Figure 1B)**. The high complementarity between the co-anchor and anchor strand forces two cholesterol molecules into close proximity. Motivated by this observation, we introduced PEG spacers to increase separation and flexibility between the DNA duplex and membrane anchor (**Figure 1A**).

**Figure 1:**
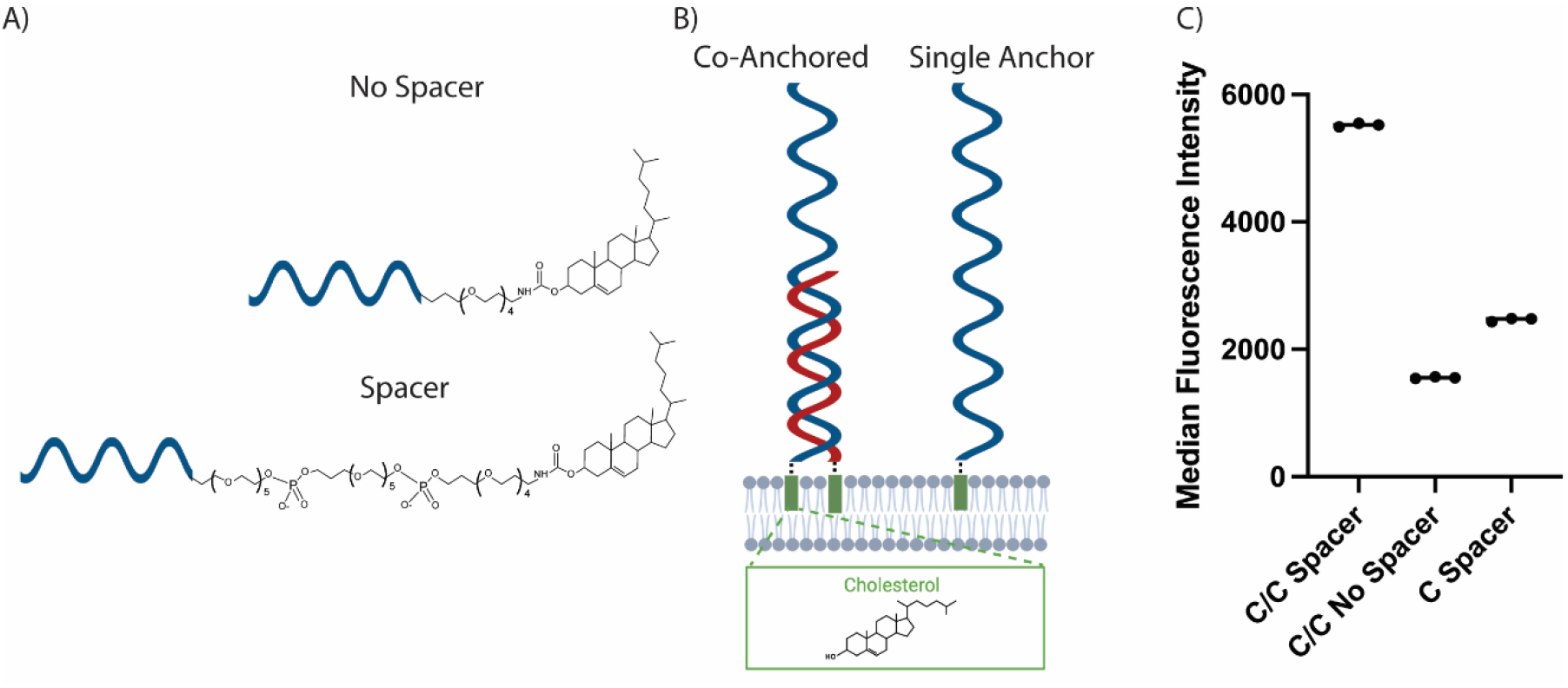
Comparisons between co-anchored and single-anchored cells with and without spacers. A) Structure of oligonucleotides with and without additional spacers between cholesterol and DNA. B) Oligonucleotide attached to a cell membrane with either a co-anchored or a single-anchored cholesterol. C) Median fluorescence intensities of different anchoring strategies on cells upon capture of fluorophore-labeled DNA.

In our initial study, we first examined whether higher DNA loading was present on the cell surface when different anchors were used. To determine, we incubated the cells with the anchor strand. After 5 minutes, we then added the co-anchoring strand. As a control, we include a spacer containing DNA without a co-anchor (C/-). The cells were rinsed to remove poorly adhering DNA. A fluorophore-labeled DNA was then introduced to the cells. Flow cytometry analysis revealed that the cholesterol/cholesterol (anchor/co-anchor; C/C) with spacers had a greater presence on the cell surface (Figure 1A). The results also indicated that the spacers consistently exhibited increased fluorescence, in contrast to the tested C/C without a spacer-anchoring strategy.

To further assess the stability of different anchoring strategies to ensure observations were not due purely to changes in DNA length, we performed an assay in which Cy3-labeled DNA was hybridized to anchor strands and then embedded on the surfaces of cells fluorescently labeled with CellTracker Deep Red (CTDR) using our cholesterol anchors/co-anchors. The anchors and co-anchors were allowed 5 minutes each to embed into the cell surface. The CTDR cells with DNA embedded on the surface were then mixed with undyed cells (**Figure S1A).** Importantly, mixed populations of CTDR cells remain distinguishable from undyed cells, allowing us to track both populations for Cy3 fluorescence (**Figure S1B**). Flow cytometry analysis of Cy3 DNA fluorescence shows that the C/C with spacers remains stable on the cell surface, as evidenced by a persistent difference in cell fluorescence (**Figure S1C**). C/C without spacers shows a decrease in the fluorescence-intensity difference between unlabeled cells and CTDR cells, such that both populations begin to exhibit similar DNA-specific fluorescence over time. Notably, single anchors show no difference in fluorescence intensity, which is expected given their limited stability in membranes.^33^

We further assessed stability by embedding the C/C with and without spacers along with C/-into cell monolayers. All anchor strands were pre-incubated with a photocaged capture strand. The photocage DNA (PC-DNA) has a region complementary to the recognition region of the anchor strand, enabling hybridization. The other PC-DNA that does not hybridize with the anchor strand features five 7-(diethylamino)-4-(hydroxymethyl)coumarin (DEACM) photocages on guanines within a 20-base-pair recognition region. We previously showed that the recognition region cannot bind to complementary DNA unless it is exposed to 425 nm light for 60 seconds (energy dose = 1.914 mJ/cm^2^). After sequentially adding anchor/co-anchors, we rinsed off the excess and exposed the cells to 425 nm light in a circular pattern. We then added Cy5-labeled DNA complementary to the photocaged regions, rinsed, and imaged to assess pattern fidelity and brightness.

Cell surfaces functionalized with the anchor and co-anchor constructs containing spacers exhibited noticeably brighter Cy3 fluorescence upon light activation, indicating greater availability on the cell surface (**Figure 2A**). Similarly, treated with C/C without spacers or C/- were noticeably dimmer with more fluorescent spillover into un-patterned regions (**Figure 2B**).

**Figure 2:**
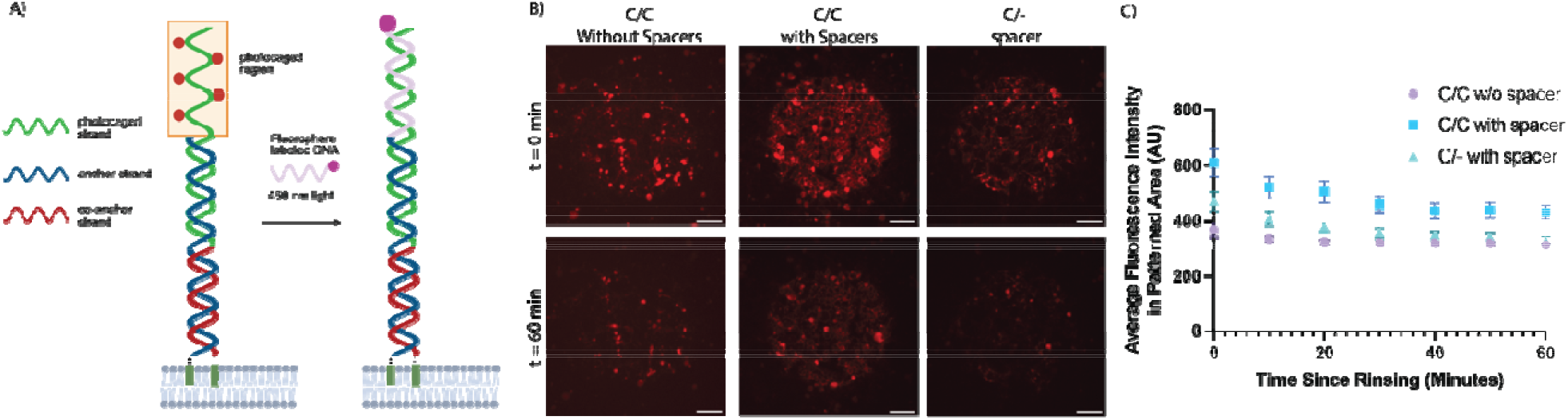
Printability and stability of DNA on cell surfaces. A) Schematic of DNA photocaging. B) Microscopy images of captured fluorophore-labeled DNA with different anchoring strategies after patterning at 0 min and 60 min post-incubation. C) Average background corrected intensities of patterned areas over time. Scale bars = 200 µm. N = 4 patterns/condition; mean +/- standard error.

Pattern stability was assessed during a 1-hour incubation at 37°C (**Figure 2C**). Fluorescence signals C/C with spacers showed greater availability across all time points. By comparison, patterns formed with spacer-free sequences or without co-anchors exhibited a progressive loss of fluorescence intensity, with visible degradation of pattern boundaries during the 1-hour incubation at room temperature in serum-free media. These results highlight that limiting dissociation is critical to achieving stability.

### Evaluating other anchoring types

For future designs, we included the PEG spacers in all the DNA we tested based on our observations. In light of our results, we synthesized an anchor with a DNA terminus containing both tocopherol and cholesterol (T-C). We hypothesized that this anchor could improve stability in the lipid membrane. We compared anchor stability using a liposomal exchange assay. For this assay, we synthesized liposomes with Cy5-labeled lipids (0.1 mol%). We incubated DNA with cholesterol only and tocopherol-cholesterol. The T-C anchors showed stronger surface interactions than the C anchors. After 5 minutes, we added a 10-fold excess of unlabeled liposomes, and the unquenching of the Cy3 signal decayed more quickly from the surface than from the more stable T-C anchors (**Figure S2**).

Based on these results, we synthesized co-anchors consisting of cholesterol (C), tocopherol (T), or tocopherol-cholesterol (T-C), again with spacers. To test the biocompatibility of our anchors/co-anchors, we performed an MTT assay. We incubated cells with all combinations of anchors/co-anchors as we did for all patterning experiments. After the fixed incubation periods we removed the media and then checked viability the following day. No change in viability was observed compared to untreated cells (**Figure S3**)

For initial printing stability studies, we first focused on whether we saw any difference in initial loading on the cell surface. For this experiment, we incubated cells with the anchor with photocaged capture strands attached for 5 minutes, then with the co-anchor. As in the previous experiments, we then exposed the patterned areas to light and incubated with a fluorophore-labeled DNA complementary to the photocaged region. This supported sustained functional DNA availability on cell surfaces during incubation at 37°C (**Figure 3, S4)**. In these experiments, tocopherol co-anchor strands consistently performed better than cholesterol-containing co-anchors over 1 h. Cholesterol anchors also showed apparent advantages over tocopherol-cholesterol anchors.

**Figure 3:**
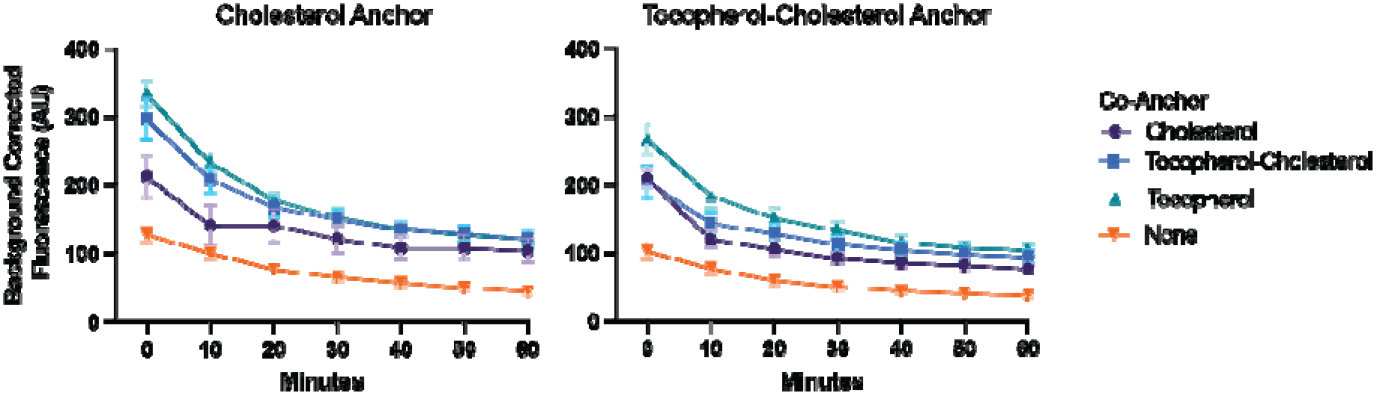
Evaluation of DNA stability on cell surface after printing. Left: Cholesterol is used as an anchor with all co-anchor combinations. Right: tocopherol-cholesterol is used as an anchor with all co-anchor combinations. N=4 patterns/condition Mean +/- standard error.

Next, we evaluated whether functional DNA availability on the cell surface was maintained over time. To evaluate the retention of accessible DNA on the cell surface, we first coated cells with anchor/co-anchor pairs and then rinsed away excess anchors/co-anchors. We then incubated the cells at room temperature and added complementary fluorophore-labeled DNA at fixed time intervals. We then assessed if we observed changes in the median fluorescence intensity compared to the initial time point. Only the single-stranded DNA anchors (T-C/- and C/-) along with the C/T-C pairing showed appreciable reductions in accessible DNA over time (**Figure 4A**).

**Figure 4:**
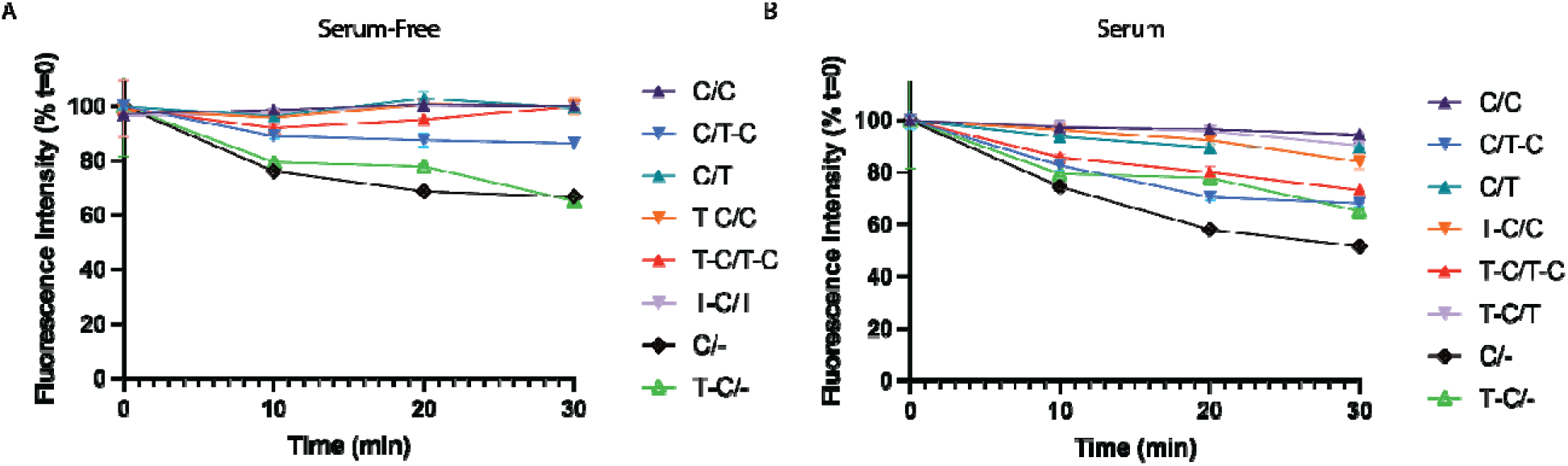
Retention of accessible DNA on cells following incubation in serum-free and serum-containing media. A) Fluorescence intensity levels after incubating cells at room temperature in serum-free media over time, followed by incubation with a fluorophore capture probe. B) Fluorescence intensity levels after incubating cells at room temperature in serum-containing media over time, followed by incubation with a fluorophore capture probe. N=3 samples for all conditions and sample types. Data presented as: mean +/- standard error.

Next, we evaluated whether this behavior changed in the presence of media. Previous studies have reported that serum-free media is required for printing. Serum contains proteins that scavenge cholesterol, tocopherol, and other lipidated molecules as they detach from the surface.^27, 30, 33^ For these studies, we incubated the cells with anchor/co-anchor DNAs in serum-free medium to prevent a large reservoir of DNA interaction before insertion into the cell surface. However, after the initial two-step incubation, we added full-serum medium followed by the fluorophore-labeled complement. For these studies, stability was highest for C/C followed by T-C/C, T-C/T, and then C/T. The T-C/T-C and C/T-C showed the lowest stability, again highlighting that poor fluidity in the co-anchor limits lock-in to the surface (**Figure 4B**).

### Cell-Printing with Different Anchors

Based on these initial studies, we performed cell printing experiments. Here, we coated cover slips with streptavidin to capture DNA terminated with biotin. The surfaces were blocked with BSA to limit nonspecific attachment. Again, we designed the biotin-terminated DNA to be partially complementary to the capture region of a PC-DNA. We then exposed the coverslips to circular patterns of 425 nm light to uncage select regions of DNA. Cells were coated with complementary DNA to the photocaged DNA (**Figure 5A**). The linker DNA on the cell surface was held to the surface using an anchor/co-anchor consisting of either C/C, C/T, or T-C/C. These combinations showed the best resilience in previous studies.

**Figure 5:**
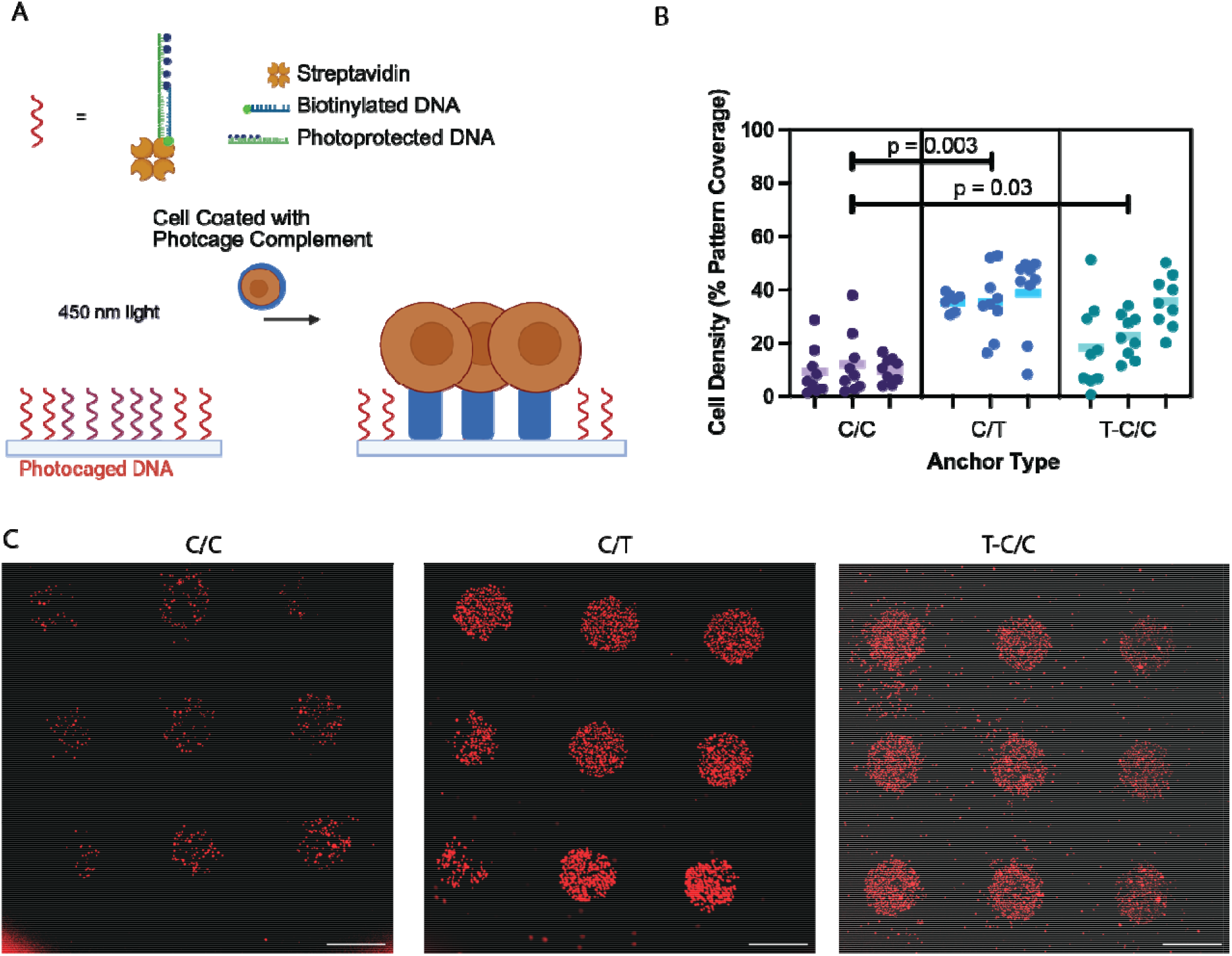
Cell printing using LD-DPC with different anchor combinations. A) Schematic of cell printing using glass coverslips coated with photocaged DNA. B) Cell printing efficiency in terms of the number of cells per pattern with different anchors. N=3 samples with 9 patterns/sample; statistics presented as Welch’s ANOVA with Tukey HSD post-hoc test. C) Micrographs of cell tracker-stained cells captured in circular patterns on a coverslip. Scale bars = 500 µm.

Cells were incubated on the surface for 5 minutes, then rinsed with serum-free medium at room temperature. The cells were then imaged to confirm placement in patterns (**Figure S5)**. Quantification of cell density as percent coverage in each patterned area showed that the C/T and T-C/C anchors performed better than the C/C anchor (**Figure 5B,C**). We also evaluated the fraction of cells within the patterns compared to the total number of cells in the field of view. For each anchor/co-anchor the median fraction was .8 or higher (**Figure S6**) indicating strong association to the patterned areas. To assess cell viability, we assessed 2 patterned substrates with C/C anchors with live/dead staining after patterning and found that >99% of the patterned cells remained viable (**Figure S7**), highlighting strong cell survival and reinforcing our previous observations with MTT assays.

We speculate that these observations may be related to differences in membrane organization. Tocopherol and cholesterol have been reported to exhibit distinct membrane partitioning behaviors, which could influence the lateral presentation of surface-bound DNA.^34–36^ While the present study does not directly measure membrane organization or diffusion, the enhanced printing performance observed with tocopherol-containing co-anchors is consistent with improved accessibility of cell-surface DNA.

As a final test, we performed a second patterning where C/T DNA anchors/co-anchors. Here, PC DNA was added to the printed cells on the surface using a two-step addition of anchor DNA pre-incubated with photocaged DNA and then co-anchor DNA (**Figure 6A**). The PC DNA was then activated on some of the cells along with areas on the coverslip not initially activated for initial cell captures. A second set of cells labeled with complementary CellTracker fluorophores was added to the surface, rinsed, and then imaged. Micrographs of the cells show that we primarily added cells to the surface in areas activated by the second light exposure, both on the coverslip and in overlapping regions of the cells (**Figure 6B, S8**). The result shows a pathway for successive, spatially controlled cell addition to a surface.

**Figure 6:**
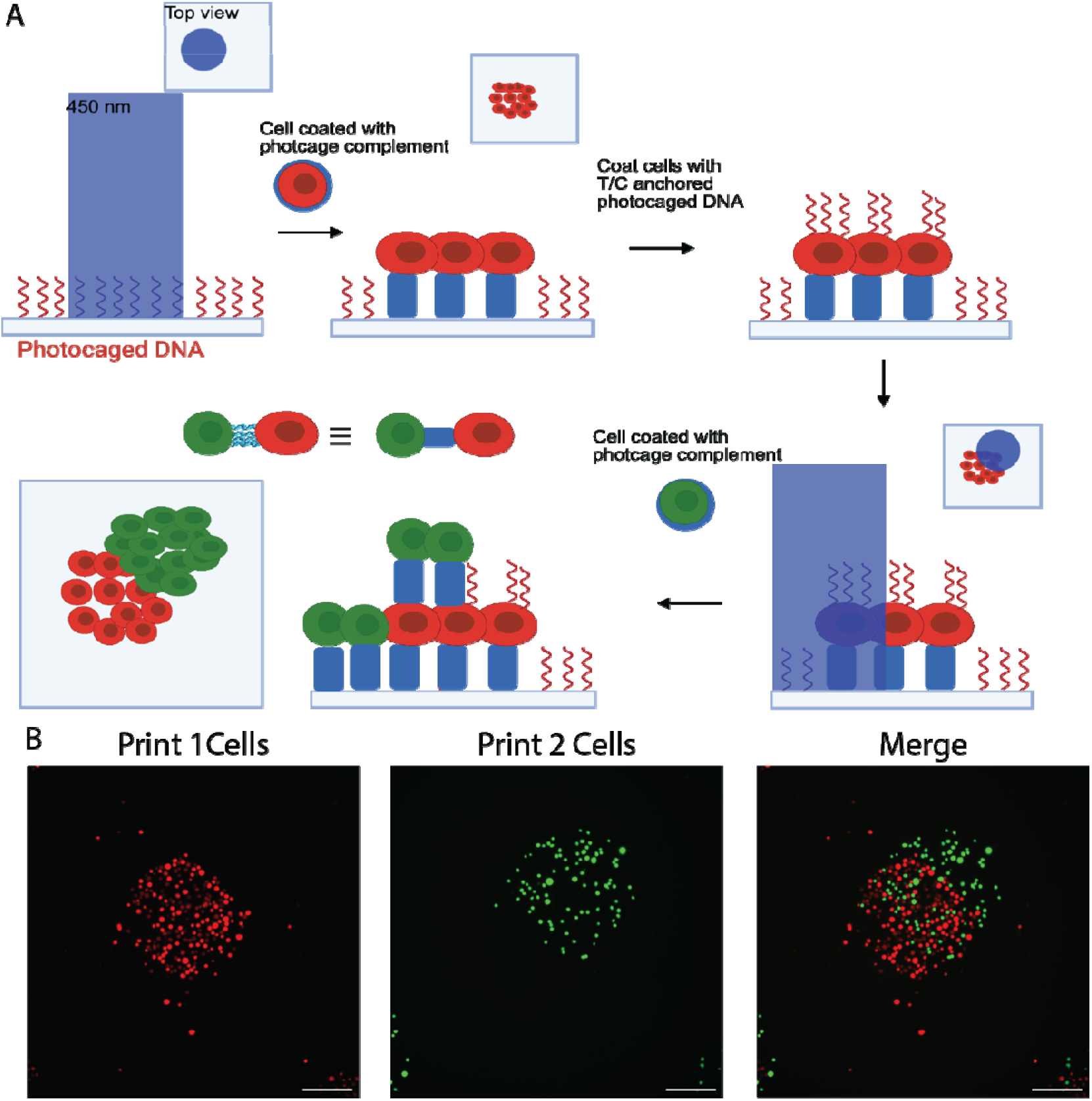
Two-layer printing. A) Schematic of a two-layer printing workflow. B) Fluorescence micrographs of cells printed in a two-layer process. Cells printed in the first print are labeled in red, and those in the second cell addition are green. Scale bars = 200 µm.

### Evaluating Phototoxicity of Patterning Light

We next evaluated whether the 425 nm light exposure used during cell patterning induced phototoxicity. We exposed cells using an inverted patterning system previously developed in our lab, calibrated to deliver the same total energy used during the standard 60s patterning protocol. Cells were exposed to 1x, 2x, 5x, or 7x the calculated energy dose (1.914 mJ/cm^2^), while unexposed cells and ethanol-treated cells served as the negative and positive controls, respectively. After exposure, cells were incubated for 24 hours and viability was quantified by an MTT assay, normalized to the unexposed control (**Figure S9**). Cells exposed to up to seven times the calculated light dose maintained viability, with no significant differences observed between any light-exposure group and the negative control. In contrast, the ethanol-treated positive control exhibited significantly reduced viability. These results indicate that the 425 nm light exposure used for cell patterning does not induce detectable phototoxicity under these conditions.

## Conclusion

Anchor and co-anchor chemistries influence the stability of molecules embedded in cell membranes, a factor that is increasingly important for light-directed DNA printing of cells (LD-DPC) and other cell-surface engineering applications. We evaluated commercially available cholesterol-, tocopherol-, and cholesterol-tocopherol-modified oligonucleotides with and without spacer elements to determine how lipid identity and molecular architecture affect functional DNA presentation at the cell surface. Our results demonstrate that both lipid identity and molecular architecture influence the availability of membrane-associated DNA. Incorporation of spacer elements enhanced DNA accessibility and printing performance, while anchor/co-anchor combinations containing tocopherol promoted efficient cell capture during printing. Together, these findings establish practical structure-function relationships that can guide the design of lipid-DNA interfaces for cell printing, cell barcoding, and programmable cellular assembly.

While the present study identifies clear relationships between anchor design and functional DNA availability, the molecular mechanisms underlying these effects, including possible contributions from membrane organization, anchor diffusion, and internalization pathways, warrant further investigation. Beyond light-directed cell printing, these findings provide practical guidance for engineering stable and accessible nucleic-acid-modified cell-material interfaces for tissue engineering, cell barcoding, and programmable multicellular assembly. Importantly, commercially available lipid-DNA chemistries maintained functional DNA presentation under room-temperature and serum-containing conditions and enabled robust single- and multilayer cell patterning. These results lower barriers to adoption of DNA-directed cell assembly approaches and provide a foundation for development of increasingly complex multicellular architectures.

## Experimental Procedures

### Safety

This work does not present any unexpected or significant hazards. Human cell lines were sourced from authorized providers and handled in compliance with institutional biosafety protocols. All chemicals were handled in accordance with standard laboratory safety procedures, with appropriate personal protective equipment used at all times.

### DNA Synthesis

All DNA monomers and reagents were purchased from Glen Research. DNA was synthesized with the Expedite synthesizer following the manufacturer’s standard protocols. We used standard bases along with the photoreactive coumarin-protected guanine (5’-Dimethoxytrityl-N2-(4-isopropylphenoxyacetyl)-O6-[[7-(diethylamino)coumarin-4-yl]-methyl]-2’-deoxyGuanosine, 3’-[(2-cyanoethyl)-(N,N-diisopropyl)]-phosphoramidite), α-Tocopherol-TEG phosphoramidite (1-Dimethoxytrityloxy-3-O-[(9-DL-α-tocopheryl)-triethyleneglycol-1-yl]-glyceryl-2-O-[(2-cyanoethyl)-(N,N,-diisopropyl)]-phosphoramidite), 5’-Cholesteryl-TEG phosphoramidite (10-O-[1-propyl-3-N-carbamoylcholesteryl]-triethyleneglycol-1-[(2-cyanoethyl)-(N,N-diisopropyl)]-phosphoramidite), and 3’-Cholesteryl-TEG CPG (1-Dimethoxytrityloxy-3-O-(N-cholesteryl-3-aminopropyl)-triethyleneglycol-glyceryl-2-O-succinoyl-long chain alkylamino-CPG). All oligonucleotides were synthesized with the dimethoxytrityl-on for purification. Oligonucleotides were deprotected using water saturated with ammonium hydroxide (28-32%; Fisher Chemicals) for 24 hours at room temperature and then dried under air (Organomation). Oligonucleotides were then purified using a Glen-Pac DNA purification column (Glen Research) following the manufacturer’s protocols. Samples were subsequently lyophilized (Labconco, -105 °C) and reconstituted to target dilutions in water.

### Cell Culture

NM2C5 cells (ATCC; CRL-2918) were maintained in DMEM with 4.5 g/L glucose, L-glutamine, and sodium pyruvate (Corning), supplemented with 10% fetal bovine serum (FBS) and 1% penicillin-streptomycin (PS). Cells were cultured at 37°C and 5% CO2. Cells were passaged at 80% confluency, and the media was changed every 2 to 3 days. Live-cell staining was performed using CellTracker (Invitrogen). CellTracker dye was resuspended in dimethyl sulfoxide (DMSO) to a 10 mM stock and diluted 1:1000 in complete media. This working solution was added to the cells for 45 minutes at 37°C. Cells were imaged using an Olympus BX63 microscope with the relevant filter sets. Imaging was completed on the same day for each experiment using the same microscope settings.

### Flow Cytometry of Mixing Treated and Untreated Cells

Cells treated with CellTracker Deep Red (CTDR) were functionalized with the DNA anchor and coanchor sequences. Stock solutions of DNA were prepared to a final concentration of 20 µM in 1X phosphate buffered saline (PBS; Hyclone Cytiva). The PBS contains 144 mg/L potassium phosphate monobasic, 9000 mg/L sodium chloride, and 795 mg/L sodium phosphate dibasic.

For each condition, 50,000 cells were suspended in 50 µL of serum-free media (DMEM with 1% penicillin–streptomycin). An additional 50 µL of either Sol. 1, 2, or 3 (**Table 1**) was added and incubated with the cells for 5 minutes. Subsequently, 100 µL of PBS or Sol. 9, 10, 11, or 12 (**Table 1**) was added, and the mixture was incubated for another 5 minutes. Following functionalization, the cells were transferred to a 96-well plate and mixed 1:1 with an equal number of undyed, untreated cells. Flow cytometry was performed exactly 5 minutes after mixing treated and untreated cells and repeated every 5 minutes for a total duration of 30 minutes. For each time point, the median Cy3 fluorescence of dyed and undyed cell populations was quantified to determine the relative amount of Cy3-labeled DNA on each group of cells.

**Table 1:** Anchor and Co-anchor Solution Components.

|  | Solution Name | Components (Sequences from Table S1) |
|---|---|---|
| **1** | C Anchor 1 | Seq. 4 + Seq. 12 |
| **2** | T-C Anchor 1 | Seq. 5 + Seq. 12 |
| **3** | C Anchor 1 without spacers | Seq. 10 + Seq. 12 |
| **4** | C Anchor 2 | Seq. 4 + Seq. 2 |
| **5** | T-C Anchor 2 | Seq. 5 + Seq. 2 |
| **6** | C Anchor 2 without spacers | Seq. 10 + Seq. 2 |
| **7** | C Anchor 3 | Seq. 4 + Seq. 9 |
| **8** | T-C Anchor 3 | Seq. 5 + Seq. 9 |
| **9** | C Co-anchor | Seq. 6 |
| **10** | T Co-anchor | Seq. 7 |
| **11** | T-C Co-anchor | Seq. 8 |
| **12** | C Co-anchor without spacers | Seq. 11 |

### Flow Cytometry for Testing Membrane Availability

For each condition, 150,000 NM2C5 cells were suspended in 50 µL of serum-free media. An additional 50 µL of 20 µM anchor DNA (Sequence 4, 5, or 10; **Table S1**) in PBS was added, and the cells were incubated for 5 minutes. Then, 100 µL of PBS with or without 20 µM coanchor DNA (Sequence 6, 7, 8, or 11; **Table S1**) in PBS was added to the cells for an additional 5 minutes. The cells were then centrifuged at 300 g for 5 minutes, and the supernatant was removed. They were resuspended with 50 µL of serum-free or serum-containing media. After time points of 0, 10, 20, and 30 minutes, 50 µL of 10 µM complementary DNA (Sequence 12; **Table S1**) in PBS was added to the corresponding cell suspension and incubated for 5 minutes. They were then centrifuged to remove excess fluorescent DNA, resuspended in a 1% paraformaldehyde solution in PBS, and incubated for 10 minutes. Finally, they were centrifuged, resuspended in PBS, and split into 3 separate wells of a 96-well plate. Flow cytometry was performed to determine the relative amount of Cy3-labeled complementary DNA attached to the anchor DNA.

### Patterning on Cell Monolayers

NM2C5 cells were grown to confluence in 24-well plates. The cells were rinsed twice with serum-free media and then incubated with Sol. 4, 5, or 6 (**Table 1**). The plate was protected from light and left for 5 min. The wells were rinsed twice with serum-free media, incubated with PBS or Sol. 9, 10, 11, or 12 (**Table 1**), protected from light and left for 5 min. The wells were then rinsed twice and covered with serum-free media.

The cells were then exposed to a 450 nm LED (ThorLabs, laser power of 1445 mW) in a circle pattern for 60s using a digital micromirror device (DMD; Mightex) attached to a microscope (BX63; Olympus). The cells were incubated with 1 µM Cy3-labeled complementary DNA (Sequence 3, **Table S1**) for 1 minute, then rinsed 3 times with serum-free medium. Stability of patterns was assessed by incubating at 37℃ for 1 hour in serum-free media and the patterns were imaged every 10 minutes using a fluorescence microscope (BX63; Olympus) with the same exposure conditions.

Quantification of pattern brightness was assessed using ImageJ. Intensity profiles were obtained for 1000 µm × 50 µm rectangles centered at the midpoint of the circle. The intensity at each point was averaged for 4 different pattern areas.

### Multilayer Cell Patterning

22 mm coverslips (Fisher Scientific) were treated with plasma oxygen (PE-25, Plasma Etch, Inc.) for 5 minutes. The coverslips were then coated with 0.01% poly-L-lysine (Sigma-Aldrich) for 30 minutes, followed by incubation with 200 nM streptavidin (Invitrogen) in 1X PBS for 1 hour. Rinse the coverslips three times with PBS. Then, the streptavidin-coated coverslips were covered with a solution of 10 µM biotinylated DNA (Sequence 1, **Table S1**) and 10 µM photocaged DNA with 5 coumarin-protected bases (Sequence 2, **Table S1**) for 10 minutes. Protect the coverslips from light after this step. Rinse the coverslips three times with PBS and cover with PBS. We then exposed the surfaces to 450 nm light in a circular pattern using a 10X objective for 60 seconds via a DMD. The coverslips were then covered with 1% bovine serum albumin (BSA; Fisher Bioreagents) for 30 minutes and rinsed 3 times with PBS.

Cells treated with CellTracker Orange (CTO), CellTracker Deep Red (CTDR), or CellTracker Green (CTG) were then functionalized with the DNA anchor and coanchor sequences. 1.5 million cells were suspended in 50 µL of serum-free media and 20 µM of 50 µL of Sol. 7 or 8 (**Table 1**) was added to the cells and left on a rotary mixer for 5 minutes. Then, 20 µM of 100 µL of Sol. 9 or 10 (**Table 1**) was added to the cells and left for an additional 5 minutes on the rotary mixer. The cells were then centrifuged at 500g for 5 minutes then resuspended in 100 µL of serum-free media.

The CTO or CTDR cell suspensions were added directly to the patterned coverslips after the PBS was removed. They were left for 5 minutes, then gently rinsed 3 times with serum-free medium. A solution of 250 µL of serum-free media and 250 µL of 20 µM of Sol. 4 or 5 (**Table 1**) was added directly to the patterned area and left for 5 minutes on the shaker. Then, 500 µL of 10 µM Sol. 9 or 10 (**Table 1**) was added directly to the coverslip for 5 minutes on the shaker, then rinsed gently three times with serum-free media. The wells were covered with serum-free medium, and the cell pattern circles were then exposed to 450 nm light using a DMD with a 10X objective for 60 seconds, in a partially overlapping pattern. CTG, CTO or CTDR cell suspensions were added to the wells after the media was removed and left for 5 minutes. The wells were gently rinsed three times with serum-free media and then covered with serum-free media.

Images were acquired using relevant filter sets and a 4x objective so that all patterns were captured within a single field of view. Images were analyzed with ImageJ. To correct for uneven illumination, images were processed using a Gaussian filter. The patterned area was selected using a circular region with a 490 µm radius. Images were then thresholded until all cells were visible, and particles smaller than 40µm^2^ were excluded from analysis. Two measurements were obtained: first, the percentage of the patterned area covered by cells, and the cell-covered area inside the patterned regions compared to outside the patterned regions.

### Viability Assays

Optical power was measured using a power meter with the same circular pattern and wavelength used for printing, for a mean of 0.0319 mW/cm^2^. The calculated energy dose was found to be 1.914 mJ/cm^2^. Cells were seeded in a 96-well plate at 20,000 cells/well, allowed to recover overnight, and exposed to 1x, 2x, 5x, or 7x the calculated energy dose. Untreated wells were used as live controls, and wells treated with ethanol for 15 minutes were as negative controls. After all conditions were incubated for 24 hours, cell viability was measured using a Sigma Aldrich MTT Cell Growth Assay kit, following the manufacturer’s protocol. Data were normalized to untreated cells, which were set to 100% viability.

For viability studies with DNA anchors, cells were seeded in a 96-well plate at 20,000 cells/well and left to recover overnight. To evaluate DNA compatibility, equal parts of serum-free media and 5 µM anchor DNA were added to each well and incubated for 5 minutes, then 5 µM co-anchor DNA or PBS was added to double the volume for an additional 5 minutes. Wells were rinsed with PBS, then replenished with complete media for 24 hours. Live and dead controls, cell viability measurement, normalization, and analysis were performed as described for viability following light exposure.

To assess the viability of cells following printing, patterned cells were stained using (Biotium, Viability/Cytotoxicityn Assay Kit for Animal Live &Dead Cells) and incubated for 15 minutes at room temperature. Cells were then imaged using appropriate filters with a 4x objective. Live and dead cells were counted in ImageJ, both within the patterned region and across the entire field of view.

## CRediT authorship contribution statement

Danielle Herubin: Writing – review & editing, Writing – original draft, Visualization, Validation, Methodology, Investigation, Conceptualization. Katelyn Mathis: Writing – review & editing, Writing – original draft, Visualization, Validation, Project administration, Methodology, Investigation, Conceptualization. Lamees Abu-Suleiman: Writing – review & editing, Writing – original draft, Validation, Methodology, Investigation. Jenna Li: Writing – review & editing, Investigation. Brian Meckes: Writing – original draft, Writing – review & editing, Visualization, Supervision, Resources, Project Administration, Funding Acquisition, Conceptualization.

## Declaration of competing interest

The authors declare no competing financial interests.

## Supporting Information

The submission includes supporting information, including additional figures and data.

## Acknowledgments

Research reported in this publication was supported by the National Institute of General Medical Sciences of the National Institutes of Health under award number R35GM150577 (B.M.). The content is solely the authors’ responsibility and does not necessarily represent the official views of the National Institutes of Health.

## Data availability

Data will be made available on request.

## Notes

### Competing Interest Statement

The authors have declared no competing interest.

## References

(1) Xie, R.; Pal, V.; Yu, Y.; Lu, X.; Gao, M.; Liang, S.; Huang, M.; Peng, W.; Ozbolat, I. T. A comprehensive review on 3D tissue models: Biofabrication technologie s and preclinical applications. Biomaterials 2024, 304, 122408. DOI: 10.1016/j.biomaterials.2023.122408 DOI.org (Crossref).

(2) Nam, K.-H.; Smith, A. S. T.; Lone, S.; Kwon, S.; Kim, D.-H. Biomimetic 3D Tissue Models for Advanced High-Throughput Drug Screenin g. SLAS Technology 2015, 20 (3), 201–215. DOI: 10.1177/2211068214557813 DOI.org (Crossref).

(3) Vunjak-Novakovic, G.; Scadden, David T. Biomimetic Platforms for Human Stem Cell Research. Cell Stem Cell 2011, 8 (3), 252–261. DOI: 10.1016/j.stem.2011.02.014 DOI.org (Crossref).

(4) Cabral, K. A.; Patterson, D. M.; Scheideler, O. J.; Cole, R.; Abate, A. R.; Schaffer, D. V.; Sohn, L. L.; Gartner, Z. J. Simple, Affordable, and Modular Patterning of Cells using DNA. Journal of Visualized Experiments 2021, (168), 61937. DOI: 10.3791/61937 DOI.org (Crossref).

(5) Chen, Z.; Fu, P.; Wang, K. DNA-programmed cell assembly: from cells, tissues to organoids. Frontiers in Bioengineering and Biotechnology 2025, 13, 1716071. DOI: 10.3389/fbioe.2025.1716071 DOI.org (Crossref).

(6) Todhunter, M. E.; Jee, N. Y.; Hughes, A. J.; Coyle, M. C.; Cerchiari, A.; Farlow, J.; Garbe, J. C.; LaBarge, M. A.; Desai, T. A.; Gartner, Z. J. Programmed synthesis of three-dimensional tissues. Nature Methods 2015, 12 (10), 975–981. DOI: 10.1038/nmeth.3553 DOI.org (Crossref).

(7) Mathis, K.; Chan, C. T.; Meckes, B. Controlling cell interactions with DNA directed assembly. Advanced healthcare materials 2024, 13 (32), 2402876.

(8) Gartner, Z. J.; Bertozzi, C. R. Programmed assembly of 3-dimensional microtissues with defined cellular connectivity. Proceedings of the National Academy of Sciences 2009, 106 (12), 4606–4610.

(9) Porter, C. M.; Qian, G. C.; Grindel, S. H.; Hughes, A. J. Highly parallel production of designer organoids by mosaic patterning of progenitors. Cell systems 2024, 15 (7), 649–661.e649.

(10) Selden, N. S.; Todhunter, M. E.; Jee, N. Y.; Liu, J. S.; Broaders, K. E.; Gartner, Z. J. Chemically Programmed Cell Adhesion with Membrane-Anchored Oligonucleo tides. Journal of the American Chemical Society 2012, 134 (2), 765–768. DOI: 10.1021/ja2080949 DOI.org (Crossref).

(11) Mathis, K.; Kohon, A. I.; Black, S.; Meckes, B. Light-Controlled Cell–Cell Assembly Using Photocaged Oligonucleotides. ACS Materials Au 2023, 3 (4), 386–393. DOI: 10.1021/acsmaterialsau.3c00020 DOI.org (Crossref).

(12) Lusic, H.; Young, D. D.; Lively, M. O.; Deiters, A. Photochemical DNA activation. Organic letters 2007, 9 (10), 1903–1906.

(13) Fichte, M. A.; Weyel, X. M.; Junek, S.; Schäfer, F.; Herbivo, C.; Goeldner, M.; Specht, A.; Wachtveitl, J.; Heckel, A. Three-Dimensional Control of DNA Hybridization by Orthogonal Two-Color Two-Photon Uncaging. Angewandte Chemie 2016, 128 (31), 9094–9098.

(14) Menge, C.; Heckel, A. Coumarin-caged dG for improved wavelength-selective uncaging of DNA. Organic letters 2011, 13 (17), 4620–4623.

(15) Mathis, K.; Gaddam, S.; Koneru, R.; Sunkavalli, N.; Wang, C.; Patel, M.; Kohon, A. I.; Meckes, B. Multifunctional hydrogels with spatially controlled light activation with photocaged oligonucleotides. Cell Reports Physical Science 2024, 5 (5).

(16) Herubin, D.; Yang, A.; Gaddam, S.; Mathis, K.; Meckes, B. Protocol for the synthesis and activation of hydrogels with photocaged oligonucleotides. STAR protocols 2025, 6 (1).

(17) Mathis, K.; Kohon, A. I.; Monroy, N.; Abu-Suleiman, L.; Yang, A.; Meckes, B. Low-Cost Open Platform Digital Light Printer (OP-DLP) for 96-Well Format Hydrogel Printing and Localized Light-Activation. ACS Biomaterials Science & Engineering 2025, 12 (1), 689–699.

(18) Hsiao, S. C.; Shum, B. J.; Onoe, H.; Douglas, E. S.; Gartner, Z. J.; Mathies, R. A.; Bertozzi, C. R.; Francis, M. B. Direct Cell Surface Modification with DNA for the Capture of Primary C ells and the Investigation of Myotube Formation on Defined Patterns. Langmuir 2009, 25 (12), 6985–6991. DOI: 10.1021/la900150n DOI.org (Crossref).

(19) Roy, S.; Cha, J. N.; Goodwin, A. P. Nongenetic Bioconjugation Strategies for Modifying Cell Membranes and Membrane Proteins: A Review. Bioconjugate Chemistry 2020, 31 (11), 2465–2475. DOI: 10.1021/acs.bioconjchem.0c00529.

(20) Li, Z.; Su, T.; Yang, Y.; Zhao, H. Construction of Multicellular Neural Tissue Using Three-Dimensional Pr inting Technology: Cell Interaction. Tissue Engineering Part B: Reviews 2025, ten.teb.2024.0323. DOI: 10.1089/ten.teb.2024.0323 DOI.org (Crossref).

(21) Ho, Y.-H.; Liao, Y.; Liao, L.; Mao, T.; Guan, Y.; Xu, R. Advances of Cell Printing Technology in Organoid Engineering. Tissue Engineering Part B: Reviews 2025, ten.teb.2025.0048. DOI: 10.1089/ten.teb.2025.0048 DOI.org (Crossref).

(22) Vogel, K.; Glettenberg, M.; Schroeder, H.; Niemeyer, C. M. DNA-modification of eukaryotic cells. Small 2013, 9 (2), 255–262.

(23) Li, J.; Wang, J.; Chen, L.; Dong, Y.; Chen, H.; Nie, G.; Li, F. Self-assembly of DNA molecules at bio-interfaces and their emerging applications for biomedicines. Nano Research 2023, 16 (12), 13014–13028.

(24) Hsiao, S. C.; Crow, A. K.; Lam, W. A.; Bertozzi, C. R.; Fletcher, D. A.; Francis, M. B. DNA-coated AFM cantilevers for the investigation of cell adhesion and the patterning of live cells. Angewandte Chemie 2008, 120 (44), 8601–8605.

(25) Ali, A. A.; Bagheri, Y.; You, M. Digging into the biophysical features of cell membranes with lipid-DNA conjugates. Quarterly Reviews of Biophysics 2022, 55, e5. DOI: 10.1017/S003358352200004X DOI.org (Crossref).

(26) Weber, R. J.; Liang, S. I.; Selden, N. S.; Desai, T. A.; Gartner, Z. J. Efficient Targeting of Fatty-Acid Modified Oligonucleotides to Live Ce ll Membranes through Stepwise Assembly. Biomacromolecules 2014, 15 (12), 4621–4626. DOI: 10.1021/bm501467h DOI.org (Crossref).

(27) Zhao, B.; Tian, Q.; Bagheri, Y.; You, M. Lipid–oligonucleotide conjugates for simple and efficient cell membran e engineering and bioanalysis. Current Opinion in Biomedical Engineering 2020, 13, 76–83. DOI: 10.1016/j.cobme.2019.12.006 DOI.org (Crossref).

(28) Li, X.; Feng, K.; Li, L.; Yang, L.; Pan, X.; Yazd, H. S.; Cui, C.; Li, J.; Moroz, L.; Sun, Y.;, et al. Lipid–oligonucleotide conjugates for bioapplications. National Science Review 2020, 7 (12), 1933–1953. DOI: 10.1093/nsr/nwaa161 DOI.org (Crossref).

(29) Scheideler, O. J.; Yang, C.; Kozminsky, M.; Mosher, K. I.; Falcón-Banchs, R.; Ciminelli, E. C.; Bremer, A. W.; Chern, S. A.; Schaffer, D. V.; Sohn, L. L. Recapitulating complex biological signaling environments using a multiplexed, DNA-patterning approach. Science advances 2020, 6 (12), eaay5696.

(30) McGinnis, C. S.; Patterson, D. M.; Winkler, J.; Conrad, D. N.; Hein, M. Y.; Srivastava, V.; Hu, J. L.; Murrow, L. M.; Weissman, J. S.; Werb, Z. MULTI-seq: sample multiplexing for single-cell RNA sequencing using lipid-tagged indices. Nature methods 2019, 16 (7), 619–626.

(31) Bagheri, Y.; Chedid, S.; Shafiei, F.; Zhao, B.; You, M. A quantitative assessment of the dynamic modification of lipid–DNA pro bes on live cell membranes. Chemical Science 2019, 10 (48), 11030–11040. DOI: 10.1039/C9SC04251B DOI.org (Crossref).

(32) Hu, C.; Liu, S.; Huang, G.; Yang, F.; Li, L.; Zhang, C.; Shao, S.; Deng, X.; Liu, Q. Close Proximity of Cholesterol Anchors in Membrane Induces the Dissociation of Amphiphilic DNA Strand from Membrane Surface. ChemBioChem 2025, 26 (1), e202400748.

(33) Meckes, B.; Banga, R. J.; Nguyen, S. T.; Mirkin, C. A. Enhancing the Stability and Immunomodulatory Activity of Liposomal Spherical Nucleic Acids through Lipid-Tail DNA Modifications. Small 2018, 14 (5), 1702909. DOI: 10.1002/smll.201702909 DOI.org (Crossref).

(34) Levental, I.; Levental, K. R.; Heberle, F. A. Lipid rafts: controversies resolved, mysteries remain. Trends in cell biology 2020, 30 (5), 341–353.

(35) Pike, L. J. Lipid rafts: bringing order to chaos. Journal of lipid research 2003, 44 (4), 655–667.

(36) Canner, S. W.; Kelley, P.; Phillips, A. Q.; Feller, S. E.; Wassall, S. R. α-Tocopherol and a Polyunsaturated Phospholipid Prefer Each Other’s Company in Mixed Membranes with Raft-Forming Sphingomyelin and Cholesterol: MD Simulations. The Journal of Physical Chemistry B 2025, 129 (40), 10353–10363.

