## Supplemental Material for "Lipid Anchor Chemistry and Spacer Architecture Influence DNA Availability and Light-Directed Cell Printing"

**Table S1: DNA Sequences,** D = DEACM caged dG-CE, S = Spacer 18, X* = phosphorothioate backbone

|  | **Name** | **5’ Modification** | **Sequence** | **3’ Modification** |
| --- | --- | --- | --- | --- |
| **1** | Biotin Surface Linker |  | ACAACAACAAGCAACAACAATTTTTTTTTT | Biotin |
| **2** | Photoprotected Adaptor (PP) |  | T*T*GTTGTTGCTTGTTGTTGTTTTTTTTTTTTTTTTTTADAADAADAACGAADAADA*A* |  |
| **3** | Complement to PP |  | T*T*CTTCTTCGTTCTTCTTC*T* | Cy3 |
| **4** | C Anchor |  | A*C*AACAACAAGCAACAACAATTTTTTTTTTGTAAGCATCCAGCTGTCAC*T*SS | Cholesterol |
| **5** | CT Anchor |  | A*C*AACAACAAGCAACAACAATTTTTTTTTTGTAAGCATCCAGCTGTCAC*T*SS | Tocopherol-Cholesterol |
| **6** | T Coanchor | Tocopherol | SSA*G*TGACAGCTGGATGCTTA*C* |  |
| **7** | C Coanchor | Cholesterol | SSA*G*TGACAGCTGGATGCTTA*C* |  |
| **8** | CT Coanchor | Cholesterol-Tocopherol | SSA*G*TGACAGCTGGATGCTTA*C* |  |
| **9** | Linker between anchor and PP |  | T*T*GTTGTTGCTTGTTGTTGTTTTTTTTTTTTTTTTTTTTCTTCTTCGTTCTTCTTC*T* |  |
| **10** | C Anchor w/o Spacers |  | A*C*AACAACAAGCAACAACAAGTAAGCATCCAGCTGTCAC*T* | Cholesterol |
| **11** | C Coanchor w/o Spacers | Cholesterol | A*G*TGACAGCTGGATGCTTA*C* |  |
| **12** | Complement to Anchors |  | T*T*GTTGTTGCTTGTTGTTGT*T* | Cy3 |
| **13** | Complement to Anchors for FRET | Cy3 | ST*T*GTTGTTGCTTGTTGTTGT*T* |  |

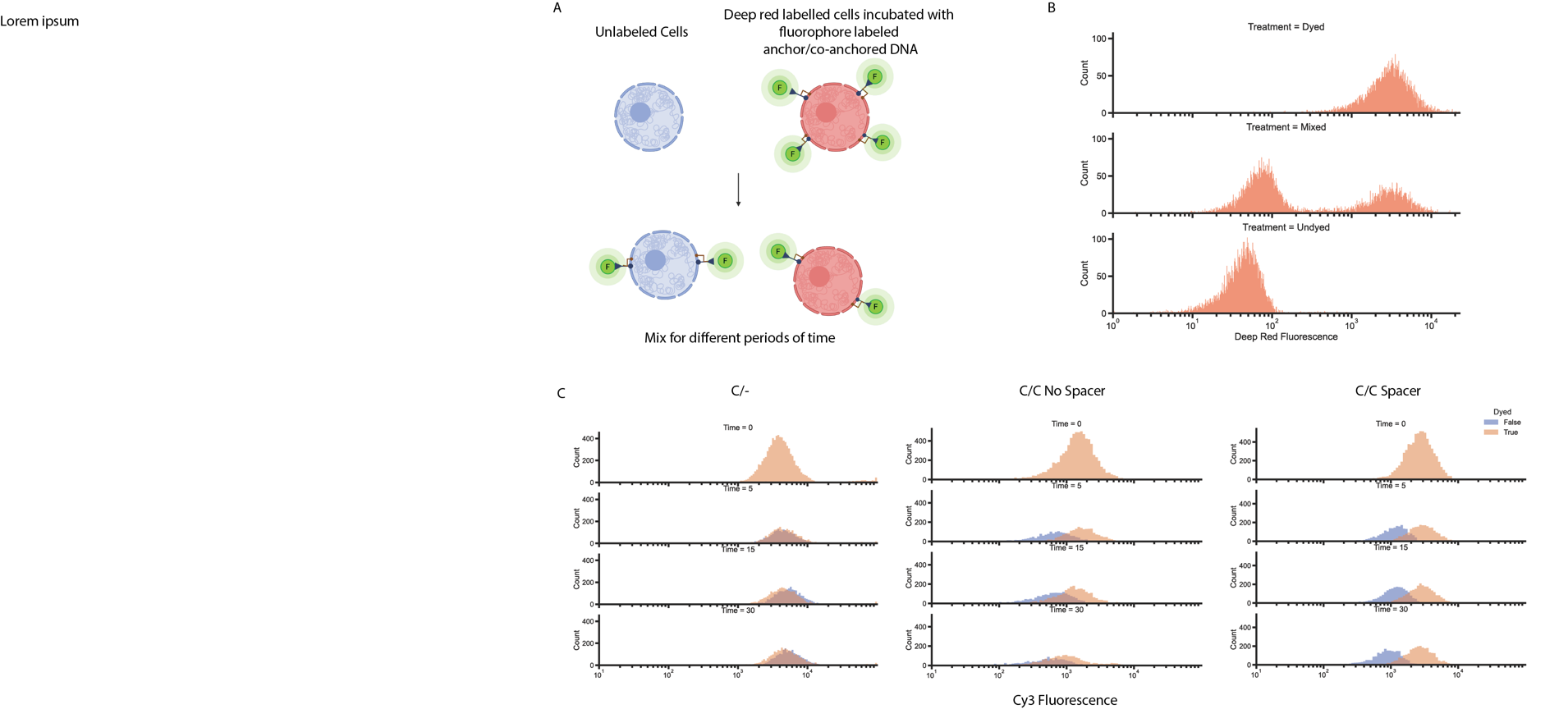

**Figure S1: Stability of anchors on cells surfaces.** A) Schematic of the experimental design where CellTracker Far Red (CTFR) cells have Cy3-labeled DNA embedded on their surface and are mixed with unlabeled cells. B) Flow cytometry of CTFR-labeled cells, unlabelled cells, and a mixed population for CTRF fluorescence. C) Flow cytometry of mixed cell populations with DNA on the surface showing the Cy3 fluorescence of the unlabeled and CTFR-labeled cells for C/-, C/C without, and C/C with spacer.

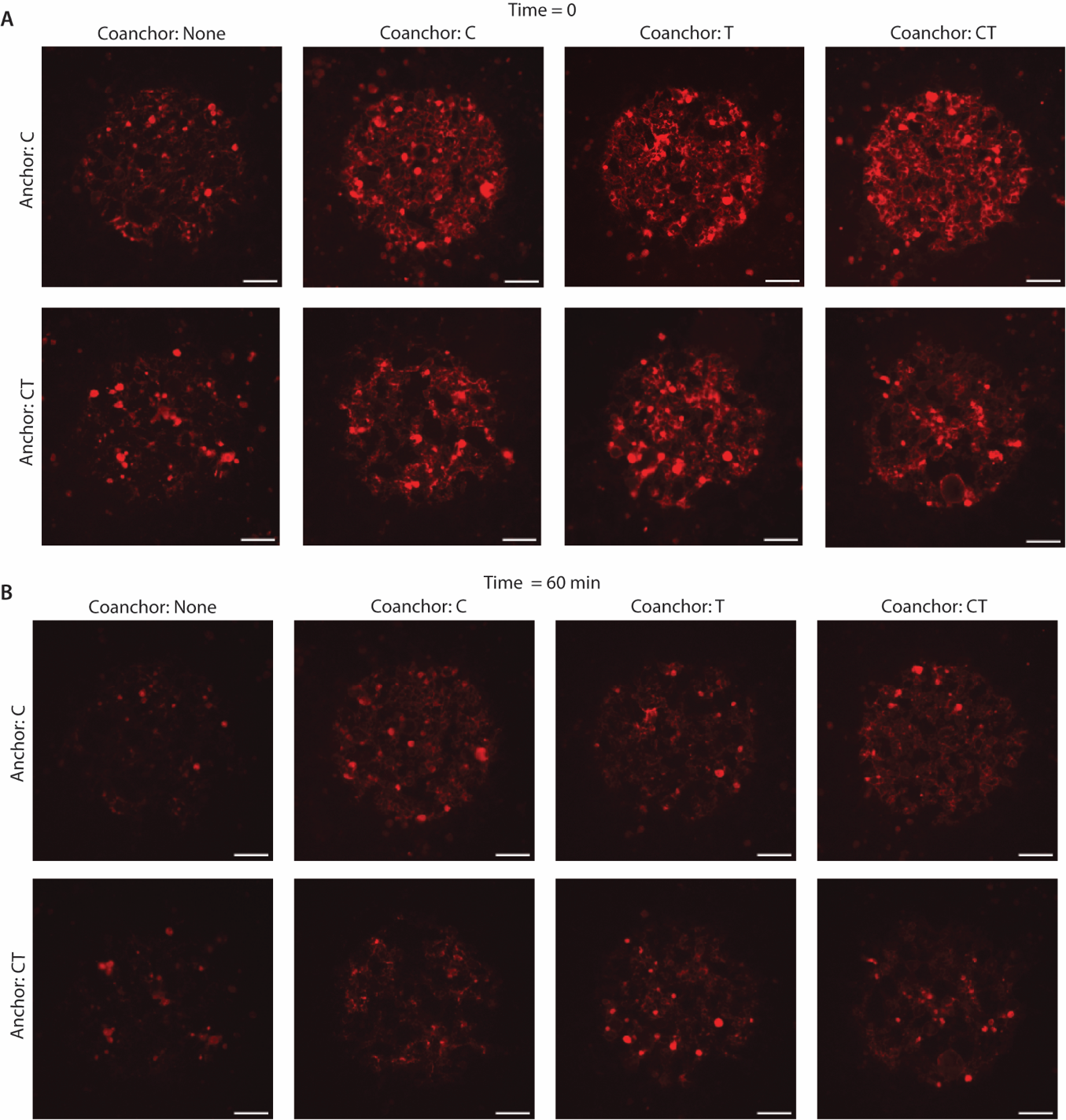

**Figure S2: Fluorescence micrographs of fluorophore DNA captured by de-caged DNA in circular patterns with different anchors**.

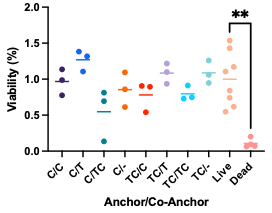

**Figure S3: MTT assay evaluating viability of cells after incubating with different anchoring/co-anchoring combinations.** **p<0.01; ANOVA with Dunnett’s comparison to the untreated live control.

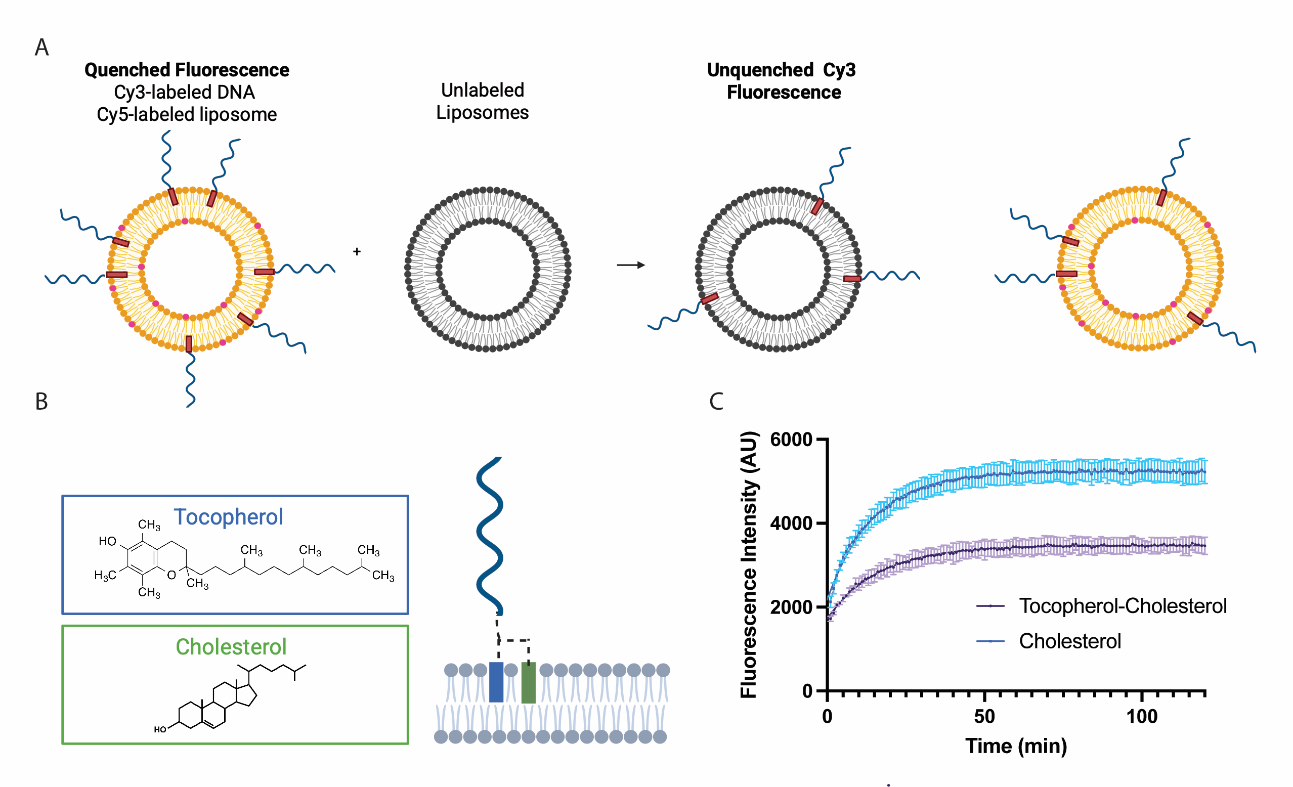

**Figure S4: Comparison between cholesterol anchoring and tocopherol-cholesterol anchoring.** A) Schematic process employed to test the exchange of anchored DNA between liposomes. B) Biochemical make-up of the tocopherol-cholesterol co-anchor molecule. C) Fluorescence dequenching due to DNA exchange between liposomes over time. N=3 for each condition; mean +/- SEM.

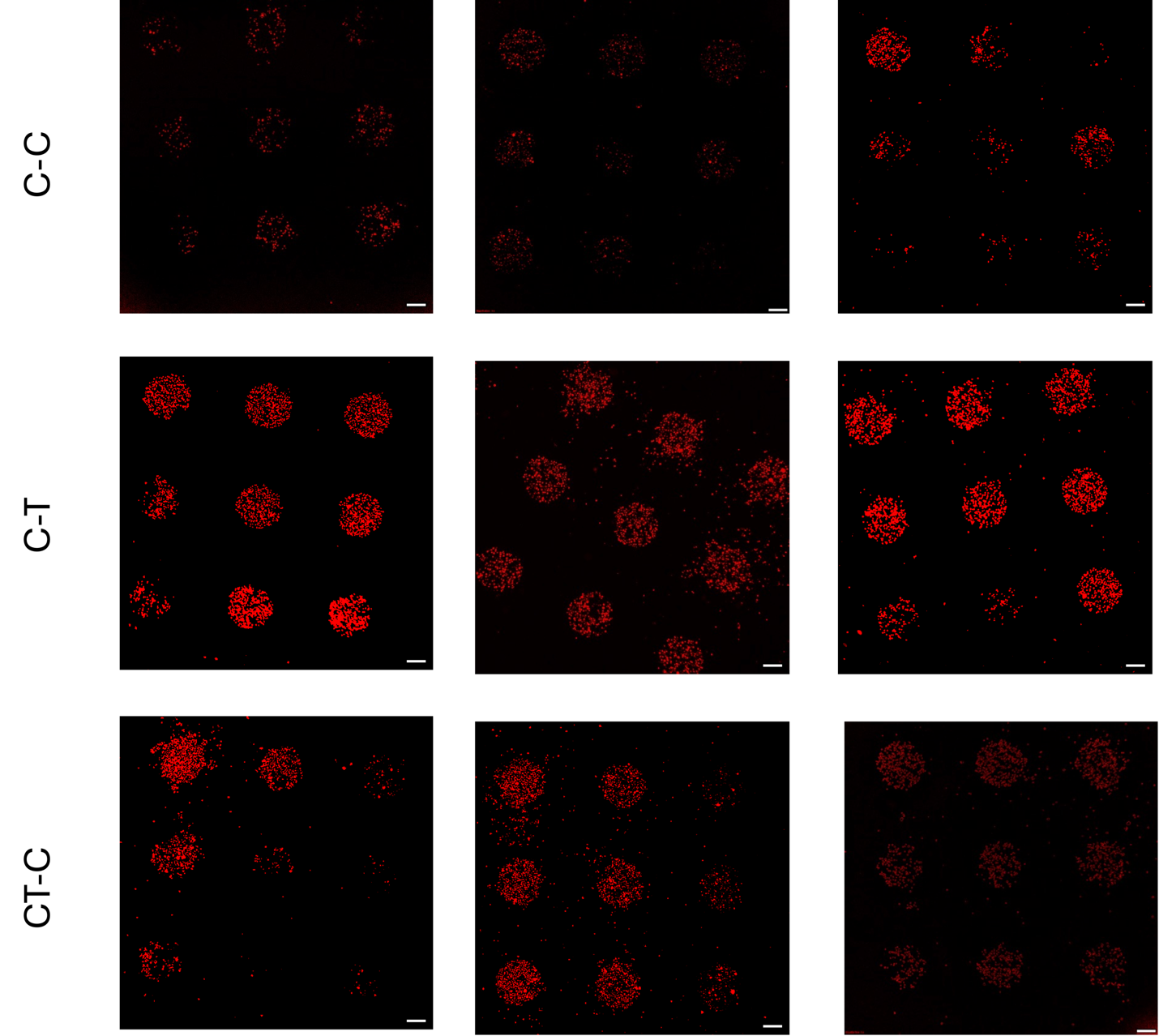

**Figure S5: Micrographs of Cell Printing Replicates. Scale bars = 200 µm.**

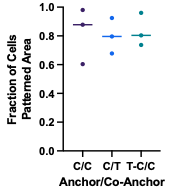

**Figure S6: Fraction of cells patterned in patterns compared to the total number of cells on the substrate within the field of view (no significant difference ANOVA).**

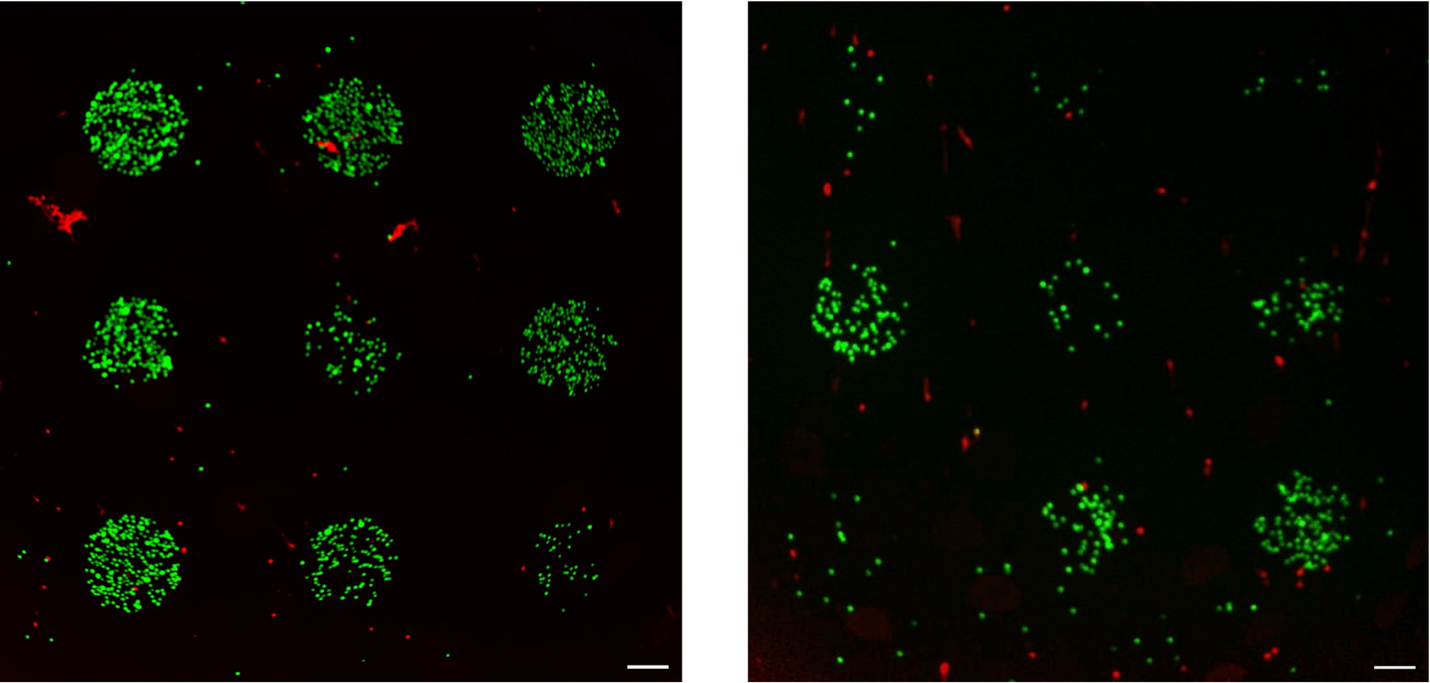

**Figure S7: Micrographs of Live/Dead stained cells patterned with C/C anchor/co-anchors. Scale bars = 200 µm.**

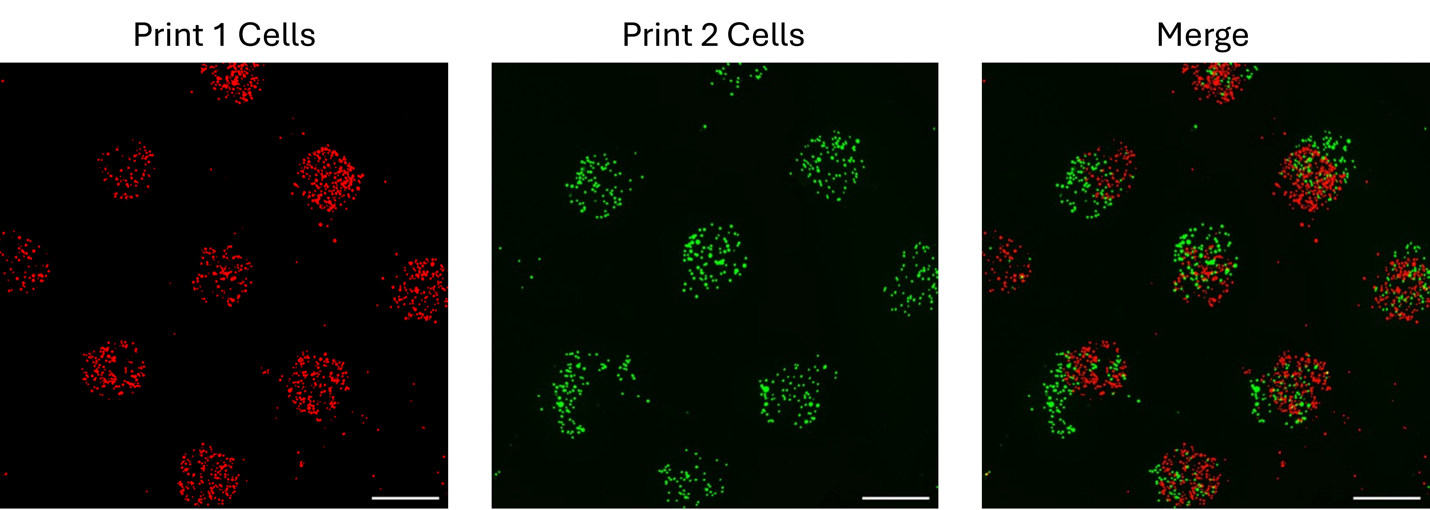

**Figure S8: Micrographs of Multilayer Cell Printing Field View. Scale bars = 500 µm.**

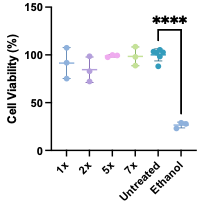

**Figure S9: MTT assays assessing viability after exposure to light equal to 1, 2, 5, and 7x doses of light used in uncaging experiments. ******p<0.0001, ANOVA with Dunnett’s test for comparing treatments to untreated controls.
